# The locus coeruleus mediates behavioral flexibility

**DOI:** 10.1101/2022.09.01.506286

**Authors:** Jim McBurney-Lin, Hongdian Yang

## Abstract

Behavioral flexibility refers to the ability to adjust behavioral strategies in response to changing environmental contingencies. A major hypothesis in the field posits that the activity of neurons in the locus coeruleus (LC) plays an important role in mediating behavioral flexibility. To test this hypothesis, we developed a novel context-dependent bilateral tactile detection task where mice responded to left and right whisker deflections in a rule-dependent manner and exhibited varying degrees of flexible switching behavior. Recording from optogenetically-tagged neurons in the LC during task performance revealed a prominent graded correlation between baseline LC activity and behavioral flexibility, where higher baseline activity following a rule change was associated with faster behavioral switching to the new rule. Increasing baseline LC activity with optogenetic activation improved task performance and accelerated task switching. Overall, our study provides strong evidence to demonstrate that LC activity mediates behavioral flexibility.

## Introduction

Behavioral flexibility, the ability to adapt goal-directed responses to changing environmental contexts and demands, is critical to the survival of organisms. For example, a pedestrian in New York should first look left to check incoming traffic before crossing a street. The same person in London would suppress this habitual response and look to the right instead. Inappropriate behavioral adaptations are observed in a broad spectrum of psychiatric disorders and aging (Uddin, 2021). Understanding the neural substrates of behavioral flexibility is a major topic of systems neuroscience research.

Several key brain structures have been implicated in supporting flexible behavioral switching (Devauges and Sara, 1990; Birrell and Brown, 2000; Lapiz and Morilak, 2006; Rich and Shapiro, 2009; Durstewitz et al., 2010; Janitzky et al., 2015; Martins and Froemke, 2015; Glennon et al., 2019; Bartolo and Averbeck, 2020), including the noradrenergic nucleus locus coeruleus (LC). A major hypothesis in the field posits that the activity of LC neurons plays a critical role in mediating behavioral flexibility (Aston-Jones et al., 1999; Aston-Jones and Cohen, 2005; Sara and Bouret, 2012). This hypothesis is primarily built upon electrophysiological evidence from non-human primates and rodents showing that LC responds to salient stimuli and that LC activity reflects behavioral states and task performance (e.g., (Aston-Jones and Bloom, 1981; Aston-Jones et al., 1994; Rajkowski et al., 1994, 2004; Usher et al., 1999; Clayton et al., 2004)). Other studies have reported that LC activity is associated with rule changes (Aston-Jones et al., 1997; Bouret and Sara, 2004; Janitzky et al., 2015; Martins and Froemke, 2015; Glennon et al., 2019; Xiang et al., 2019). However, to the best of our knowledge, it is currently unknown whether and how LC activity is linked to different degrees of behavioral flexibility. Furthermore, no prior work has directly tested this hypothesis by recording and perturbing genetically-defined noradrenergic neurons in the LC during a rule-shift task to determine causal relationships between LC activity and behavioral flexibility. Answering these questions is a critical step toward unraveling the molecular, cellular and circuit mechanisms underlying LC modulation of behavioral flexibility and cognitive functions.

To bridge these knowledge gaps and to assess the extent to which the LC contributes to flexible task switching, we developed a novel context-dependent bilateral tactile detection task in head-fixed mice, where mice were trained to respond to left and right single-whisker deflections in a rule-dependent manner and exhibited varying degrees of flexible switching behavior within individual sessions. During task performance, we recorded from optogenetically-tagged noradrenergic neurons in the LC and established a graded relationship between LC spiking activity and flexible task switching. Higher behavioral flexibility (faster switching to the new rule) was characterized by a greater increase in baseline LC activity upon the rule change compared with lower behavioral flexibility (slower switching to the new rule). Increasing baseline LC activity with optogenetic activation led to robust improvements in task switching and task performance. Together, our data provide strong evidence to support that the LC is critical in mediating flexible behavioral switching.

## Results

We developed a novel context-dependent bilateral tactile detection task to probe behavioral flexibility in head-fixed mice. There were two rules (contexts) in the task: Left Go and Right Go, and mice were trained to adapt to repeated rule changes within individual behavioral sessions (Fig. 1a-c, Methods). On each trial, one of the two whiskers (left or right C2) was stimulated. For the Left Go rule, the left whisker deflection was the Go stimulus and the right whisker deflection was the NoGo stimulus, and vice versa for the Right Go rule. Licking during a response window following whisker stimulation determined trial outcome (Fig. 1b-c). Individual sessions typically consisted of 2-3 blocks, with each block consisting of 100-200 trials. The rule of the first block (block 1) was randomly assigned, and each subsequent block had the alternate rule as the preceding block (e.g., Block 1: Left Go; Block 2: Right Go, Fig. 1a). The beginning of each block consisted of 5-10 ‘cueing trials’, where the presentation of the Go stimulus was paired with water delivery (Methods). Since block 1 did not involve a rule change, subsequent analyses were focused on the blocks following block 1, with switch performance referring to task performance (fraction correct) in these blocks. The training process took several weeks (Fig. 1d), and mice were considered well trained once their switch performance in block 2 was above 65% for two consecutive days (Chevée et al., 2021) (Methods). Mice were able to perform Left Go and Right Go blocks at similar levels (i.e., no apparent ‘handedness’, Fig. S1a), so both types of blocks were pooled in subsequent analyses.

**Figure 1.**
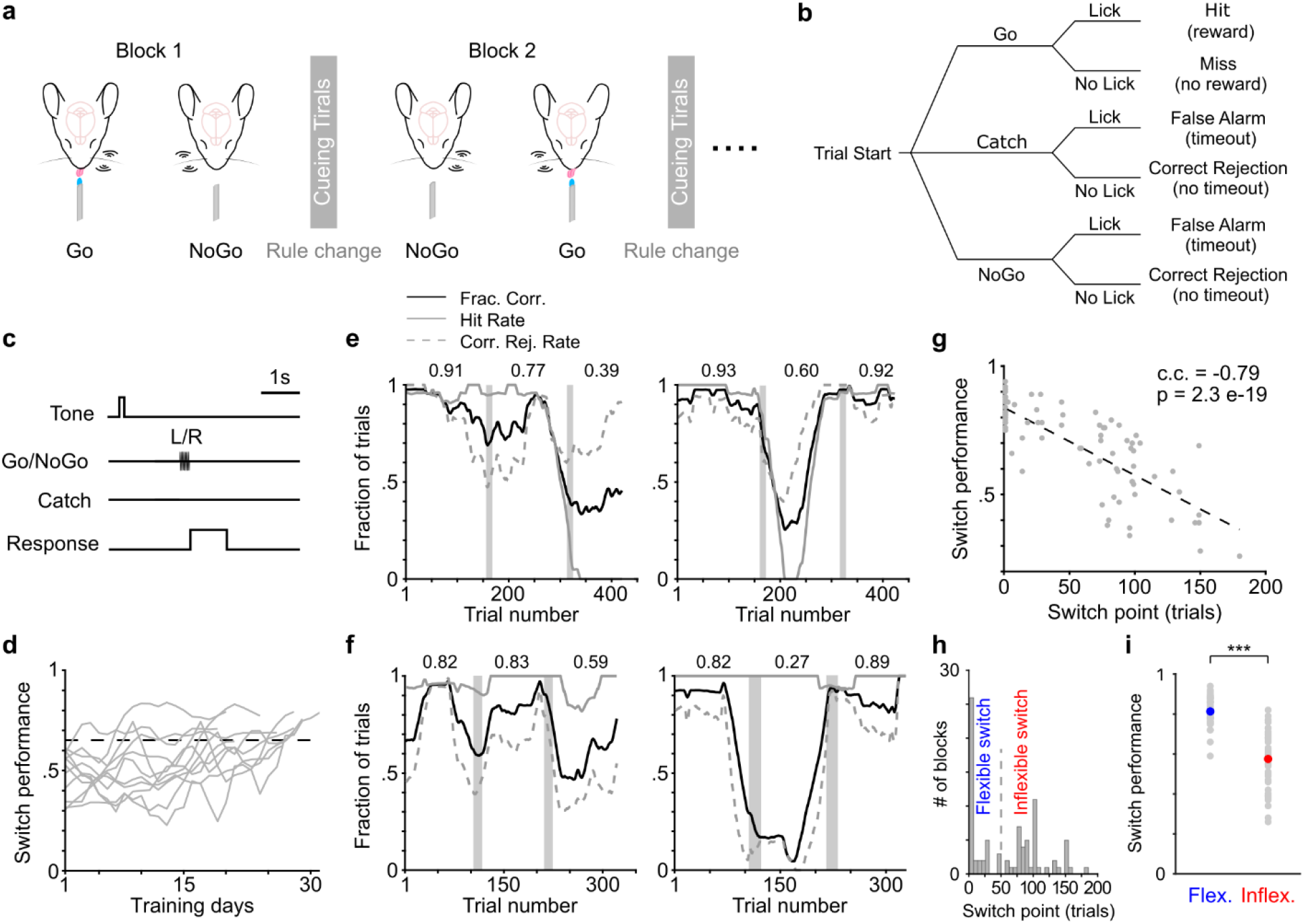
A novel context-dependent tactile detection task to probe flexible task switching. **(a-c)** Schematic of the novel context-dependent bilateral tactile detection task in head-fixed mice (a), with illustrations of trial types (b) and trial structure (c). **(d)** Switch performance (fraction correct in block 2) during training (n = 11 mice). Dotted line indicates 65% threshold. **(e, f)** Example behavioral sessions for two mice, respectively, with performance in each block indicated. Left panels illustrate higher switch performance in block 2, and right panels illustrate higher switch performance in block 3. Vertical gray bars indicate cueing trials. **(g)** The relationship between behavioral switch point and switch performance. c.c., Pearson correlation coefficient. **(h)** Histogram of behavioral switch point (84 blocks from 11 mice). We used 50 trials (dashed line) to separate flexible and inflexible blocks. **(i)** Switch performance for flexible and inflexible blocks shown in (h). Flexible (40 blocks) vs. Inflexible (44 blocks): 0.81 ± 0.011 vs 0.58 ± 0.022, p = 2.9e-12, rank sum = 2480. Gray dots represent individual blocks, blue and red dots represent mean.

We noticed that well-trained mice exhibited varying levels of switch performance following a rule change (Fig. 1e, f). To quantify the degree of flexible task switching, we defined a behavioral switch point where task performance in a 50-trial moving window surpassed a threshold of 85% (Methods). Notably, behavioral switch point exhibited a strong negative correlation with switch performance (Fig. 1g). Since behavioral switch point followed a bimodal distribution separated around trial 50, we referred to blocks with behavioral switch point below trial 50 as ‘flexible blocks’ and those above as ‘inflexible blocks’ (Fig. 1h). As expected, switch performance in flexible blocks was significantly higher than in inflexible blocks (Fig. 1i). To test whether such differences in switch performance were due to variations in motivational states (Berditchevskaia et al., 2016; Allen et al., 2019; McBurney-Lin et al., 2020), we did further analysis to show that neither flexible nor inflexible blocks were preferentially concentrated toward the beginning or the end of individual sessions (Fig. S1b). Additionally, reaction time and number of licks were not different between flexible and inflexible blocks (Fig. S1c). These results strongly suggest that motivational changes during a session cannot account for the differences in task switching between flexible and inflexible blocks. Together, our data demonstrate that mice exhibited varying degrees of flexible switching behavior in the novel context-dependent bilateral tactile detection task.

Next, we recorded spiking activity from optogenetically-tagged single noradrenergic neurons in the LC along with pupil diameter during task performance (Fig. 2a, b). Based on previous work (Aston-Jones et al., 1997; Usher et al., 1999; Aston-Jones and Cohen, 2005; Yang et al., 2021), we hypothesized that the pre-stimulus baseline LC activity was associated with the degree of behavioral flexibility. To test this, we first analyzed baseline LC activity (quantified in 1-s window preceding whisker stimulation onset) before and after the rule change in a subset of sessions where both a flexible block and an inflexible block occurred (14 blocks from 7 sessions, for paired comparison) and found that LC spiking was transiently elevated following the rule change only in the flexible blocks (Fig. 2c-e). That is, in blocks where mice more rapidly adapted their responses to the new rule, baseline LC activity was transiently elevated following the rule change (After), compared with the baseline LC activity right before the rule change (i.e., at the end of the previous block - Before, Fig. 2d-f, Methods). The changes in baseline activity upon the rule shift (ΔFiring rate: After – Before) were also higher in flexible blocks than inflexible blocks (Flexible vs. Inflexible, 0.50 ± 0.16 vs. −0.28 ± 0.16 spikes/s, p = 0.02, Fig. 2f). Similar trends held when we included additional flexible and inflexible blocks that were not from the same sessions (i.e., unpaired comparison, Fig. S2a). Importantly, the changes in baseline LC activity upon the rule shift exhibited a graded and significant negative correlation with behavioral switch point (Fig. 2g, Pearson correlation coefficient = −0.42, p = 0.0026), substantiating and extending the findings based on the binarized flexible and inflexible blocks. Licking behavior during flexible and inflexible blocks were similar (Fig. S2b), and the trend of LC activity held when we only included hit trials in the analysis (Fig. S2c, d). In addition, baseline activity was quantified prior to any possible licking events in a trial. Together, these lines of evidence strongly suggest that the observed changes in LC activity were not a direct effect of licking itself (Zagha et al., 2022).

**Figure 2.**
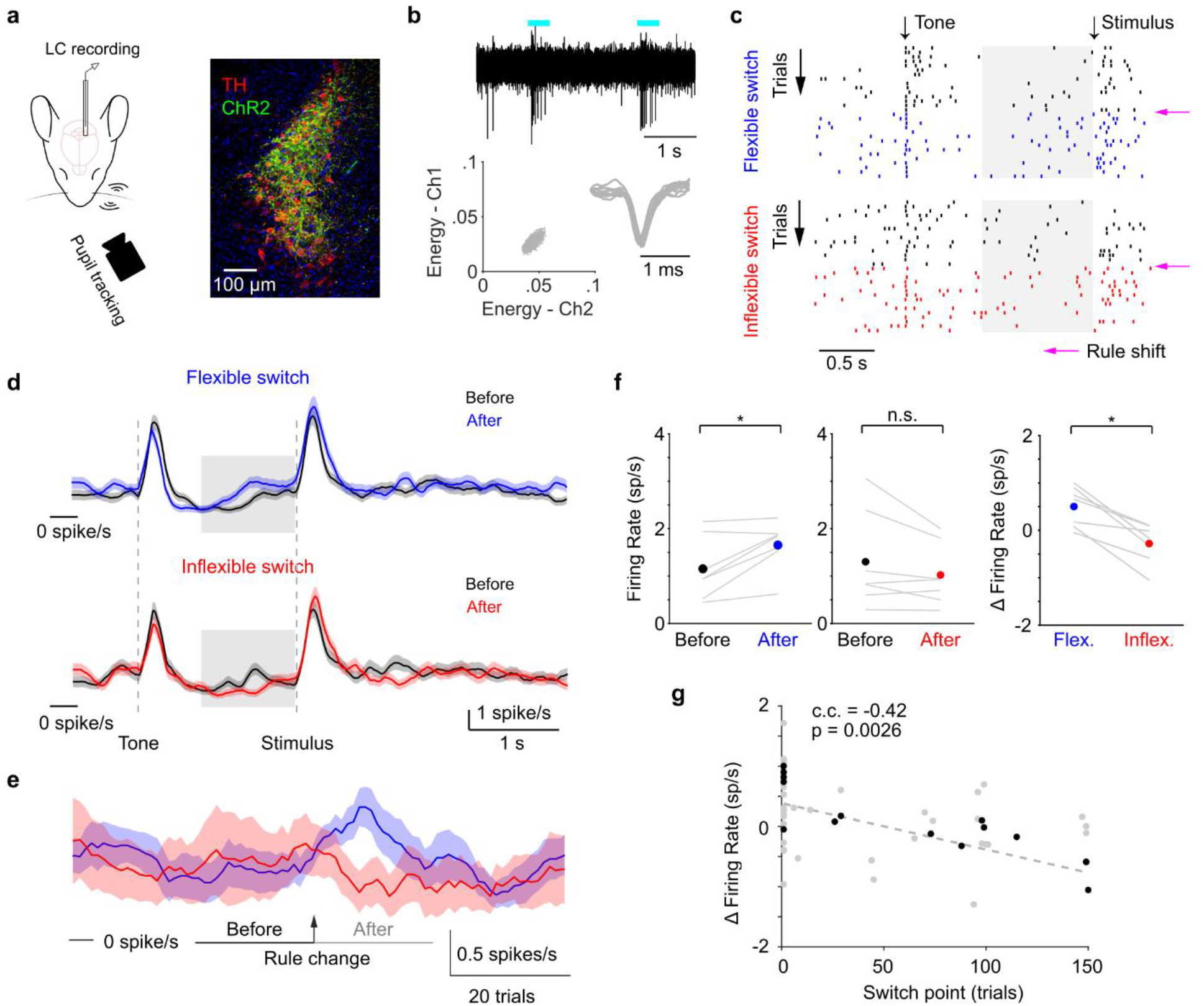
Correlating LC activity with flexible task switching. **(a)** Schematic of experimental setup during behavior (Left) and ChR2 expression in a DBH;Ai32 mouse (Right. TH: tyrosine hydroxylase). **(b)** Top: Responses of an example ChR2-expressing LC neuron to optical stimulation (cyan bars). Bottom: spike-sorting diagram and waveforms of this unit. **(c)** Example spike raster from an example LC unit in a behavior session with both a flexible (Top) and an inflexible (Bottom) block. Rows represent individual trials. Magenta arrows indicate the rule shift. Trials in the previous block are in black, and trials in the current block are in blue (flexible) or red (inflexible). Shaded gray areas represent the 1-s time window to quantify baseline activity. **(d)** Average PSTH of LC activity from flexible (Top, n = 7) and inflexible (Bottom, n = 7) switches, quantified before (last 20 trials in the previous block, Before) and after the rule change (first 20 trials in the new block, After). Shaded gray areas represent the 1-s time window to quantify baseline activity. **(e)** Average baseline LC activity in a 100-trial window centered at the rule change (arrow) for flexible (blue) and inflexible (red) switches (same blocks as in d). Horizontal bars indicate Before and After trial periods to compare baseline LC activity in d, f, g. **(f)** Baseline LC activity during the Before and After periods for flexible (Left, Before vs. After: 1.15 ± 0.25 vs. 1.65 ± 0.19 spikes/s, p = 0.03, signed rank = 1, n = 7) and inflexible switches (Middle, Before vs. After: 1.30 ± 0.39 vs. 1.02 ± 0.24 spikes/s, p = 0.16, signed rank = 23, n = 7). Changes in baseline activity (Δ Firing rate: After - Before) was higher during flexible switches than inflexible switches (Right, Flexible vs. Inflexible: 0.50 ± 0.16 vs. −0.28 ± 0.16 spikes/s, p = 0.02, signed rank = 28, n = 7). Lines represent individual paired flexible-inflexible blocks from the same session. Dots represent mean. **(g)** The relationship between behavioral switch point and the changes in baseline LC activity. Gray dots represent individual blocks (n = 49). Black dots represent the paired blocks shown in (d-f). c.c., Pearson correlation coefficient.

To further determine whether the observed changes in baseline LC activity reflected true changes in behavioral flexibility, we quantified task performance in trial blocks immediately preceding the identified flexible and inflexible blocks. Performance in blocks immediately preceding the flexible blocks (i.e., blocks with a different rule) was comparable to the performance in those flexible blocks. In contrast, task performance in blocks immediately preceding the inflexible blocks was higher than the performance in the inflexible blocks (Fig. S2e). These results strongly suggest that 1) the identified flexible blocks and the associated LC activity reflected true flexible task switching, such that mice were adapting to both rules, instead of simply following one rule; and 2) the identified inflexible blocks and the associated LC activity reflected true inability to switch to the new rule, instead of an overall lack of performance in the task. Although pupil diameter was bigger in the After period in both flexible and inflexible blocks, the changes in pupil diameter upon the rule shift (ΔPupil: After - Before) were slightly bigger in the flexible blocks than the inflexible blocks (Fig. S2f), consistent with the trend of LC activity. Overall, our data show that baseline LC activity was correlated with the degree of flexible task switching, such that higher baseline activity following the rule change was associated with a faster behavioral adaptation to the new rule.

Next, to determine the causal role of LC activity in behavioral flexibility, we optogenetically activated LC neurons during behavior. We recruited mice that had been trained for >4 weeks with switch performance consistently below the 65% threshold. Adapted from the paradigm used in a previous study (Glennon et al., 2019), optical stimulation (10 Hz, 10 mW) was delivered in a 0.5-s window prior to whisker stimulation onset in trials following the rule change (Fig. 3a, Methods). Channelrhodopsin-2 (ChR2) was expressed in the LC of both test and control groups, but the test group had the optical fiber implanted in the LC, and the control group had the optical fiber implanted away from the LC (Fig. S3a). Fiber implant was confirmed by pupil responses to optical stimulation (Fig. 3b, (Privitera et al., 2020; Megemont et al., 2022)). Switch performance of the test group noticeably improved upon LC stimulation, compared with previous sessions without stimulation (Baseline vs. Stimulation: 0.52 ± 0.02 vs. 0.69 ± 0.02 fraction correct, p = 1.6e-5), and the improvement was present in individual mice (Fig. 3c, d, f. For extended baseline periods refer to Fig. S3b, c). The behavioral effects appeared to be specific to the switching blocks as performance in the first block (block 1) was not affected (Fig. S3d). LC stimulation also accelerated task switching in the test group (Behavioral switch point, Baseline vs. Stimulation: 154 ± 8 vs. 116 ± 13 trials, p = 0.02, Fig. 3g). In contrast, optical stimulation had no effects on task switching in the control group (Fig. 3c, e-g, Fig. S3b, c). The behavioral improvement in the test group appeared to be due to an increase of correct rejection rate, but not hit rate (Fig. 3h, i, Fig. S3e, f), in line with a previous report (Glennon et al., 2019). Licking behavior was not influenced by optical stimulation thus cannot account for the increase in correct rejection rate (Fig. S3g, h). In summary, increasing baseline LC activity facilitated flexible task switching.

**Figure 3.**
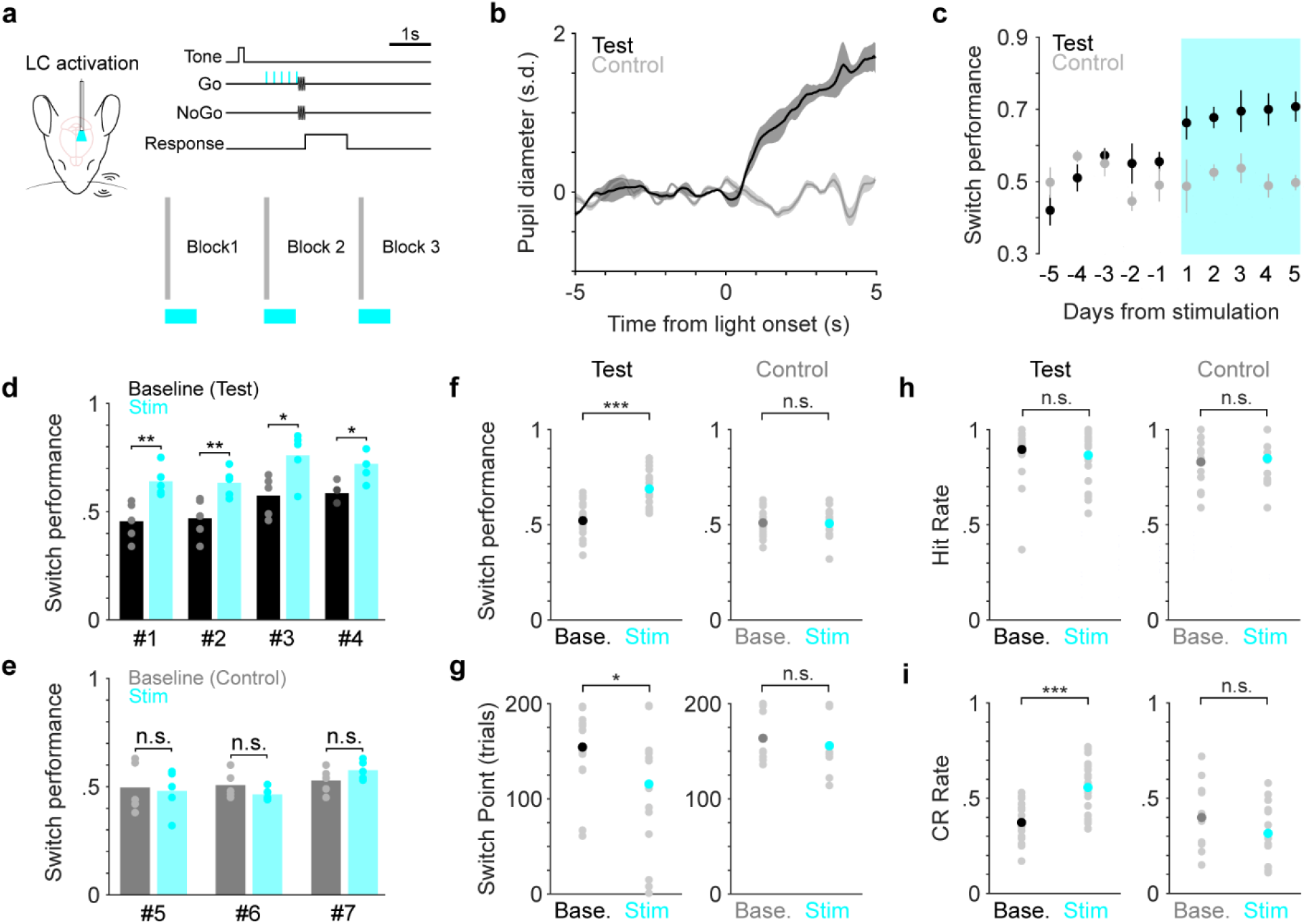
Determining the causal links between LC activity and flexible task switching. **(a)** Schematic of optogenetic LC stimulation during task performance. **(b)** Example pupil responses to optical stimulation for a test (black) and control (gray) mouse. **(c)** Group average switch performance for the test (black) and control (gray) groups during baseline (5 consecutive days prior to stimulation) and optical stimulation (5 consecutive days with stimulation, cyan) sessions. Day −1 represents the last day without stimulation. Day 1 represents the first day with stimulation. **(d)** Switch performance for individual mice in the test group (n = 4), compared between baseline (black) and stimulation sessions (cyan). Baseline vs. Stimulation, Mouse #1: 0.46 ± 0.04 vs. 0.64 ± 0.03, p = 0.008; Mouse #2: 0.47 ± 0.04 vs. 0.63 ± 0.03, p = 0.004; Mouse #3: 0.57 ± 0.04 vs. 0.76 ± 0.05, p = 0.012; Mouse #4: 0.59 ± 0.02 vs. 0.72 ± 0.03, p = 0.024. Permutation test. **(e)** Switch performance for individual mice in the control group (n = 3), compared between baseline (gray) and stimulation sessions (cyan). Baseline vs. Stimulation, Mouse #5: 0.50 ± 0.05 vs. 0.48 ± 0.05, p = 0.80; Mouse #6: 0.51 ± 0.03 vs. 0.46 ± 0.02, p = 0.24; Mouse #7: 0.53 ± 0.03 vs. 0.58 ± 0.01, p = 0.18. Permutation test. **(f)** Comparison of switch performance for test (Left) and control (Right) groups between baseline and stimulation sessions. Baseline vs. Stimulation, Test group: 0.52 ± 0.02 vs. 0.69 ± 0.02, p = 1.6e-5, rank sum = 250, n = 20; Control group: 0.51 ± 0.02 vs. 0.51 ± 0.02, p = 1.0, rank sum = 232, n = 15. Black, dark gray and cyan dots represent mean. **(g)** Comparison of behavioral switch point for test (Left) and control (Right) groups between baseline and stimulation sessions. Baseline vs. Stimulation, Test group: 154 ± 8 vs. 116 ± 13 trials, p = 0.02, rank sum = 495, n = 20; Control group: 164 ± 7 vs. 156 ± 8 trials, p = 0.23, rank sum = 262, n = 15. Black, dark gray and cyan dots represent mean. **(h)** Comparison of hit rate for test (Left) and control (Right) groups between baseline and stimulation sessions. Baseline vs. Stimulation, Test group: 0.90 ± 0.03 vs. 0.87 ± 0.03, p = 0.56, rank sum = 432, n = 20; Control group: 0.83 ± 0.03 vs. 0.85 ± 0.04, p = 0.79, rank sum = 162, n = 15. Black, dark gray and cyan dots represent mean. **(i)** Comparison of correct rejection rate for test (Left) and control (Right) groups between baseline and stimulation sessions. Baseline vs. Stimulation, Test group: 0.37 ± 0.02 vs. 0.56 ± 0.03, p = 2.4e-4, rank sum = 274, n = 20; Control group: 0.40 ± 0.04 vs. 0.32 ± 0.04, p = 0.15, rank sum = 268, n = 15. Black, dark gray and cyan dots represent mean.

## Discussion

In this work, we set out to test the hypothesis that the LC is involved in mediating behavioral flexibility. We developed a novel context-dependent bilateral tactile detection task where head-fixed mice exhibited varying degrees of flexible task switching upon a rule change within single sessions. Using this task, we found that the magnitude of baseline LC activity following the rule change was closely correlated with the degree of flexible task switching. Specifically, higher baseline activity upon the rule shift was associated with faster behavioral switching. Next, we optogenetically enhanced baseline LC activity and observed improved task performance and accelerated task switching. Overall, our study highlights the role of LC in behavioral flexibility and provides strong evidence to support that LC activity mediates flexible task switching.

Pre-stimulus baseline LC activity exhibited a relatively small (<1 spike/s) yet significant increase during flexible switching, in line with prior work showing similar magnitude of firing rate changes (commonly referred to as tonic activity) associated with behavioral states or rule shifts (e.g., Aston-Jones and Bloom, 1981; Aston-Jones et al., 1997; Usher et al., 1999; Xiang et al., 2019). Importantly, we uncovered that such activity exhibited a graded, negative relationship with the degree of behavioral flexibility. We also noted that LC neurons transiently responded to the auditory tone and whisker stimulation (commonly referred to as phasic activity in the literature), and found relatively weak relationships between such responses and task switching (Fig. S4).

Our perturbation data support recent work showing that activating the LC facilitated auditory reversal learning (Martins and Froemke, 2015; Glennon et al., 2019). One major distinction in our study is that the current task is essentially a continuous reversal task where prior to LC stimulation mice had been trained to adapt to multiple reversal stages (blocks) within single sessions, similar to the task structure of a recent study (Chevée et al., 2021). This task design allows us to assess the relationship between LC activity and rapid, ‘real-time’ behavioral switching within individual sessions. Specifically, the association between baseline LC activity and the degree of flexible task switching was transient (~20 trials, equivalent to 2-3 minutes, Fig. 2e. Also see Aston-Jones et al., 1997). Secondly, LC optical stimulation was delivered in a subset of trials (a total of ~20-30 trials per block, Methods) and the behavioral improvement was present from the first stimulation session (day 1, Fig. 3c). As a result, long-term plasticity mechanisms, such as structural synaptic changes, are unlikely to underlie such rapid associations between LC activity and behavioral switching. Nevertheless, it is worth noting that in a recent study where LC stimulation was paired with the target tone in a relatively long-term fashion, rats were found to better suppress their responses to non-target tones (Glennon et al., 2019), consistent with an increase of correct rejection rate in our task. Together, these studies suggest that LC activity facilitates behavioral flexibility across different time scales, likely through different mechanisms.

How does LC activity drive behavioral flexibility? This remains a major challenge in the field. Ample prior research has shown that noradrenergic (NA) signaling from the LC modulates neuronal responses to sensory stimuli in various sensory-related brain structures (Berridge and Waterhouse, 2003; McBurney-Lin et al., 2019). Thus, the transient increase of LC activity following the rule change may modulate neuronal responses to the new Go and/or NoGo stimulus (Devilbiss and Waterhouse, 2004; Devilbiss et al., 2006; Martins and Froemke, 2015; Rodenkirch et al., 2019) to better separate signal and noise representations. Another possibility is that LC-NA signaling facilitates the reshaping of the relevant stimulus representation to more effectively influence behavior (Ruff and Cohen, 2016, 2019). On the other hand, decades of work has established the importance of the prefrontal cortex (PFC) in behavioral flexibility (Miller and Cohen, 2001; Le Merre et al., 2021; Uddin, 2021), and LC-NA signaling heavily influences PFC functions (Arnsten and Li, 2004; Ramos and Arnsten, 2007; Arnsten et al., 2012). Transient changes in LC activity may dynamically modulate synaptic efficacy and recurrent activity in the PFC to affect top-down regulation of the relevant and irrelevant stimuli (Arnsten et al., 2012; Zagha, 2020) and to facilitate reorienting behavior (Sara and Bouret, 2012). Future experiments with simultaneous recordings from LC-NA and sensory/executive areas will elucidate how LC-NA activity modulates bottom-up processing and top-down commands to influence behavioral flexibility.

## Author contributions

J.M.L. and H.Y. planned the project and built apparatus. J.M.L. performed experiments. J.M.L. and H.Y. analyzed data and wrote manuscript.

## Acknowledgements

We thank Edward Zagha, Martin Riccomagno, Sachiko Haga-Yamanaka for discussion and comments on the manuscript. This work was supported by UCR startup, UC Regents Faculty Fellowship, Klingenstein-Simons Fellowship Awards in Neuroscience, and NIH grants (R01NS107355, R01NS112200) to H.Y..

## Data availability

Data are available from the corresponding author upon request.

## Code availability

MATLAB scripts used to analyze the data are available from the corresponding author upon request.

## Materials and Methods

All procedures were performed in accordance with protocols approved by UC Riverside Animal Care and Use Committee (AUP 20190031). Mice were DBH-Cre (B6.FVB(Cg)-Tg(Dbh-cre) KH212Gsat/Mmucd, 036778-UCD, MMRRC); Ai32 (RCL-ChR2(H134R)/EYFP, 012569, JAX), singly housed in a vivarium with a reversed light-dark cycle (12 hr/12 hr). Procedures for headpost and custom optrode microdrive implants have been described in detail previously (Yang et al., 2016, 2021). Briefly, 8-12 week old male and female mice were implanted with titanium headposts, leaving a window open above the left cerebellum for subsequent tetrode implants. Custom tetrode microdrives were made with eight tetrode wires surrounding an optic fiber (0.39 NA, 200 um core) to make extracellular recordings from opto-tagged LC neurons. The microdrive was implanted targeting the left LC. Mice were then allowed to recover for at least 72 hours before water restriction and behavior training. Once mice reached the performance threshold (65%, block 2), the microdrive was advanced at regular intervals (75 um/day) towards LC. LC neurons were identified by optogenetic tagging, tail pinch response, and post-hoc lesions. Thirty-four single-unit recordings/sessions (cluster quality measure Lratio: 0.01 ± 0.005; firing rate: 2.44 ± 0.30 spikes/s; percent ISI < 10 ms: 1.21% ± 0.42%) from six mice performing the context-dependent bilateral tactile detection task (see below) were extracted using MClust (Redish, 2014), along with synchronous recording of the left pupil. Optogenetic activation experiments were acquired from seven mice (4 test, 3 control) that had been trained for >4 weeks on the bilateral task with performance consistently below the 65% threshold. Prior to optogenetic experiments, placement of the optic fiber was assessed by pupil responses to optical stimulation under anesthesia (10-ms pulses, 10 Hz, 10 mW). At the conclusion of all experiments, electrolytic lesions were made and brains perfused with PBS, followed by 4% PFA. The brains were post-fixed in 4% PFA overnight, then cut into 100 um coronal sections and stained with anti-Tyrosine Hydroxylase (TH) antibody (Thermo-fisher OPA1-04050) and anti-EGFP antibody (Thermo-fisher A-11039).

Mice were water restricted to 1mL/day for at least seven days prior to behavioral training. Behavioral tasks were controlled via a custom-based Arduino hardware and software and acquired in WaveSurfer (https://www.janelia.org/open-science/wavesurfer), and one behavioral session was performed per day. Mice were first trained to a modified version of the Go/NoGo single-whisker detection task described previously (Yang et al., 2016; McBurney-Lin et al., 2020). In brief, mice reported the presence of a brief deflection (0.2-s, 25-Hz sinusoidal) to either the right or the left C2 whisker by licking a water port during a 1-s response window. On Go trials, stimulation of the right or the left whisker was delivered in alternating blocks (e.g., first 100 trials, stimuli are presented to the right whisker, following 100 trials to the left whisker, etc.). On NoGo trials, no whisker stimulation was delivered and mice were trained to withhold licking. Mice usually achieved >75% overall performance within 7 days. Mice were then introduced to the second stage of training, in which an identical whisker deflection was presented in NoGo trials on the contralateral side to the Go stimulus (e.g., left stimulus: Go; right stimulus: NoGo). Similar to the first stage of training, the stimuli were presented in block structures, with the Go stimulus alternating from left to right whiskers. Early during this stage of training, a single rule switch was implemented and each block consisted of 200 trials. The first block rule was randomly assigned as either left Go/right NoGo or right Go/left NoGo. As the mouse became more proficient in switching, the blocks were shortened progressively and additional 1-2 rule changes added. Therefore, as mice learned the task, more rule changes were introduced into single sessions. The mice in this study could execute between one and three rule changes in a session.

The beginning of each block consisted of a ‘cueing window’, which was a period of 5-10 consecutive Go trials where whisker stimulation was paired with water delivery, designed to facilitate adaptation to the new rule. A 0.1-s auditory cue (8 kHz, ~80 dB SPL) signaled the start of each trial, followed by a 1.5-s delay before whisker stimulation. If mice licked in this delay window, the trial was aborted and the next trial began after a 5-10 s timeout. Ambient white noise (cut off at 40 kHz, ~80 dB SPL) was played continuously to mask any potential cues that can be associated with the task. Go and NoGo trials represented 90% of all trials. Catch trials represented the remaining 10%, in which no whisker stimulation was presented, and mice were trained to withhold licking. On Go trials, the Go stimulus for the current block was delivered, and mice were expected to report its presence by licking the water port within a 1-s window immediately following stimulus cessation. Correct responses to Go stimulus presentation were qualified as ‘hit’ trials and rewarded with a water drop (~5uL). Withholding licking to Go stimulus were qualified as ‘miss’ trials. Withholding licking to NoGo stimulus and on catch trials were unrewarded and qualified as ‘correct rejection’. Licking to NoGo stimulus and on catch trials were qualified as ‘false alarm’ and punished with a 5-s timeout. If the mouse licked again within the timeout period but at least 1-s after the initial response, there was a subsequent 5-s punishment, with up to three consecutive timeouts allowed per trial. To further assist mice to suppress licking to the NoGo stimulus, a NoGo trial was designed to follow a ‘false alarm’ trial (Aruljothi et al., 2020), and up to 4 consecutive NoGo trials were allowed to occur. As a result, behavioral sessions typically consisted of more NoGo trials than Go trials (~55% vs. 45%).

Optogenetic stimulation (10-ms pulses, 10 Hz, 10 mW) was delivered using a 450 nm blue diode laser (UltraLasers, MDL-III-450-200mW) and controlled by WaveSurfer. The mating between sleeve and ferrule was covered with polymer clay to prevent light leakage. Stimulation of LC neurons was delivered on Go trials during a 0.5-s window prior to whisker stimulation onset. Video of the left pupil was acquired at 20 Hz using a Basler acA1300-200um camera and Pylon software. Pupil diameter was measured offline using DeepLabCut (Mathis et al., 2018). Electrophysiology recording, pupil tracking, and optogenetic stimulation were synchronized via a common TTL pulse train.

Switch performance (fraction correct) was quantified in blocks other than the first block (block 1) in a session. To compute the behavioral switch point in a block, task performance was first quantified using a 50-trial moving window. Behavioral switch point was defined as the beginning of the moving window within which the average task performance surpassed 85% threshold (Fig. 2 results held when using other thresholds between 75% and 85%, data not shown). If this criterion was never met, i.e., task performance in the 50-trial moving window never reached 85% threshold within the block, switch point was set as the total number of trials in that block. For Fig. 1d, switch performance was quantified in block 2 as early training sessions only had 2 blocks. For Fig. 1g-i, 84 blocks from 11 mice were included.

For Fig. 2, baseline LC activity was quantified in a 1-s window prior to whisker stimulation onset. 20 trials before the rule change (i.e., last 20 trials in the previous block) were considered as the Before period, and 20 trials after the rule change were considered as the After period. For Fig. 2d, the average firing rate in the Before and After periods was smoothed using a 150-ms window. Figure 2e quantified the 10-trial averaged baseline firing rate for ±50 trials from the rule change. The change (Δ) in baseline firing rate was calculated as baseline LC activity in the Before period subtracted from the After period (After - Before; Fig. 2f, g). For Fig. 2g, 49 blocks from 6 mice were presented, a subset of the sessions as in Fig. 1g-i.

In Fig. 3, optogenetic stimulation began at the beginning of each block and ended 50 trials after the cueing trials. Stimulation (10-ms pulse train at 10 Hz and 10 mW) was delivered on Go trials only, starting at 0.5 s before whisker stimulation onset and ending at the onset. Switch performance and switch point were quantified in block 2 as majority of the sessions had 2 blocks. 5 consecutive baseline sessions (no stimulation) and 5 consecutive stimulation sessions from each mouse were included for analysis (1 session per day). For Fig. 3d, e, analyses were performed for individual mice separately. For Fig. 3 f-i, sessions were pooled from all mice in each condition (i.e., 20 sessions from the test group (4 mice) and 15 sessions from the control group (3 mice) in each condition).

Data were reported as mean ± s.e.m. unless otherwise noted. We did not use statistical methods to predetermine sample sizes. Sample sizes are similar to those reported in the field. We assigned mice to experimental groups arbitrarily, without randomization or blinding. Statistical tests were two-tailed Wilcoxon signed rank (paired) or rank sum (unpaired) unless otherwise noted.

