## Supplemental Figures for "The locus coeruleus mediates behavioral flexibility"

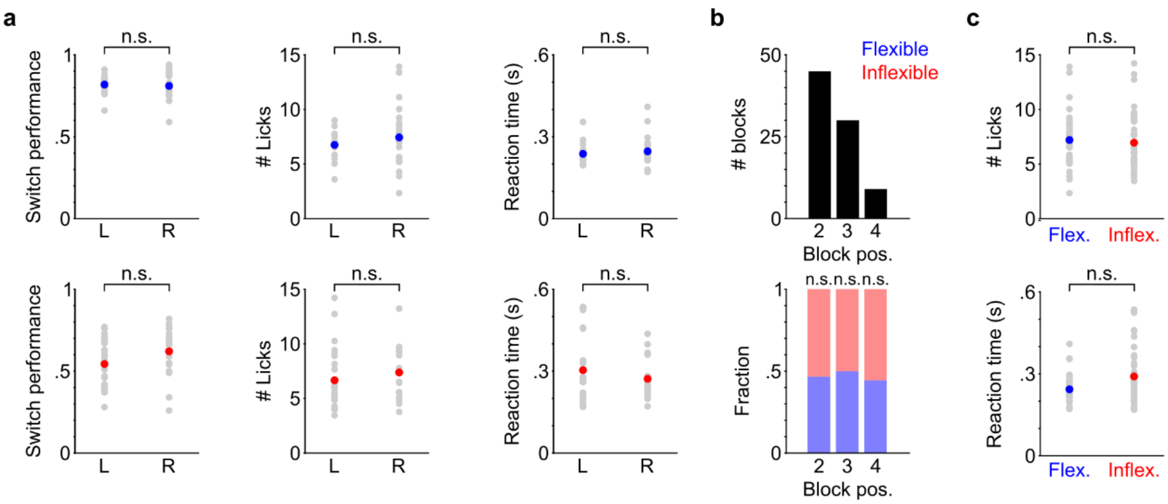


**Figure S1.**

**(a)** Behavioral variables quantified for Left Go blocks (L) and Right Go blocks (R) in flexible (Blue, Top: L, n = 14; R, n = 26) and inflexible (Red, Bottom: L, n = 26; R, n = 18) switches. Flexible: switch performance, L vs. R: 0.82 ± 0.02 vs. 0.81± 0.01, p = 0.71, rank sum = 301; Number of licks, L vs. R: 6.76 ± 0.38 vs. 7.44 ± 0.53, p = 0.42, rank sum = 258; Reaction time, L vs. R: 0.24 ± 0.01 vs. 0.25 ± 0.01 s, p = 0.41, rank sum = 257. Inflexible switches: switch performance, L vs. R: 0.54 ± 0.03 vs. 0.62 ± 0.04, p = 0.095, rank sum = 515; Number of licks, L vs. R: 6.66 ± 0.53 vs. 7.38 ± 0.58, p = 0.25, rank sum = 536; Reaction time, L vs. R: 0.30 ± 0.02 vs. 0.27 ± 0.02 s, p = 0.59, rank sum = 608. Gray dots represent individual blocks, red and blue dots represent mean. Number of licks and reaction time were quantified in hit trials. Reaction time was calculated as the latency from whisker stimulation onset to the time of the first lick.

**(b)** The distribution of block positions within a session for the blocks shown in Fig. 1g-i (Top, n = 84), and the proportion of flexible (blue) and inflexible (red) blocks in each position within a session (Bottom). Block 2: Flexible vs. Inflexible: 0.47 vs. 0.53, p = 0.53, t-stat = 0.40; block 3: Flexible vs. Inflexible: 0.50 vs. 0.50, p = 1, t-stat = 0; block 4: Flexible vs. Inflexible: 0.44 vs. 0.56, p = 0.64, t-stat = 0.22, Chi-squared test.

**(c)** Comparison of lick responses between flexible and inflexible blocks (40 vs. 44 blocks). Number of licks, Flexible vs. Inflexible: 7.20 ± 0.37 vs. 6.96 ± 0.39, p = 0.38, rank sum = 1800; Reaction time, Flexible vs. Inflexible: 0.24 ± 0.01 vs. 0.29 ± 0.02 s, p = 0.056, rank sum = 1486. Gray dots represent individual blocks, blue and red dots represent mean.

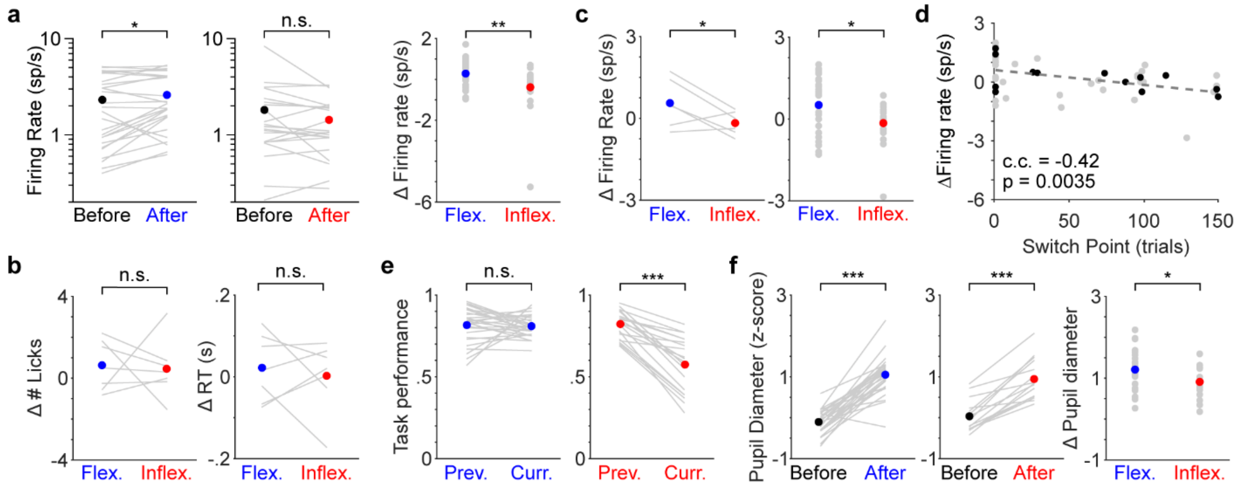


**Figure S2.**

**(a)** Baseline LC activity during the Before and After periods for flexible (Left: Before vs. After: 2.31 ± 0.31 vs. 2.60 ± 0.30 spikes/s, p = 0.03, signed rank = 109.5, n = 28) and inflexible (Middle: Before vs. After: 1.81 ± 0.38 vs. 1.43 ± 0.20 spikes/s, p = 0.10, signed rank = 163, n = 21) switches. Changes in baseline activity (ΔFiring rate: After - Before) were higher during flexible switches than inflexible switches (Right: Flexible vs. Inflexible: 0.29 ± 0.12 vs. -0.38 ± 0.26 spikes/s, p = 0.0074, rank sum = 833).

**(b)** Changes in number of licks (Left: Δ# licks, After – Before, Flexible vs. Inflexible: 0.79 ± 0.52 vs. 0.56 ± 0.62, p = 0.96, t-stat = -0.05, n = 6) and reaction time (Right: ΔRT, Flexible vs. Inflexible: 0.03 ± 0.04 vs. 0.03 ± 0.02 s, p = 0.83, t-stat = 0.22, n = 6) for the paired flexible-inflexible blocks shown in Fig. 2d-f.

**(c)** Changes in baseline LC activity for the paired flexible-inflexible blocks (n = 6) as in Fig. 2d-f (Left, Flexible vs. Inflexible: 0.56 ± 0.36 vs. -0.18 ± 0.18 spikes/s, p = 0.041, t-stat = 2.16, two-tailed t-test) and all blocks (n = 47) as in Fig. S2a (Right, Flexible vs. Inflexible: 0.51 ± 0.20 vs. -0.15 ± 0.17 spikes/s, p = 0.037, rank sum = 722) quantified in hit trials only. For b and c, there were no hit trials during the Before period of 2 blocks from the original full dataset (n = 49), and they were removed from analyses. 1 block was included in the original paired analysis (n = 7), so the associated flexible-inflexible pair was removed from analyses.

**(d)** The relationship between behavioral switch point and the changes in baseline LC activity quantified in hit trials only as shown in (c). Conventions are as in Fig. 2g.

**(e)** Left: Comparison of task performance quantified in blocks immediately preceding the flexible blocks (Previous) and quantified in the flexible blocks (Current). Previous vs. Current: 0.82 ± 0.02 vs. 0.81 ± 0.01, p = 0.29, signed rank = 157, n = 28. Right: Comparison of task performance quantified in blocks immediately preceding the inflexible blocks (Previous) and quantified in the inflexible blocks (Current). Previous vs. Current: 0.82 ± 0.02 vs. 0.57 ± 0.03, p = 5.9e-5, signed rank = 0, n = 21.

**(f)** Pupil diameter during the Before and After periods for flexible (Left, Before vs. After: -0.10 ± 0.06 vs. 1.05 ± 0.09 s.d., p = 4.7e-6, signed rank = 404, n = 28) and inflexible (Middle, Before vs. After: 0.045 ± 0.08 vs. 0.93 ± 0.13 s.d., p = 1.8e-4, signed rank = 188, n = 19) switches. Changes in pupil diameter (ΔPupil diameter: After - Before) was higher during flexible switches than inflexible switches (Right, Flexible vs. Inflexible: 1.15 ± 0.10 vs. 0.88 ± 0.11 s.d., p = 0.044, rank sum = 751). 2 inflexible blocks with poor pupil tracking were excluded from this analysis. Pupil diameter was calculated as the average pupil diameter during a baseline pupil window. The baseline pupil window was defined as a 1-s window starting 1.5 s after the start of the baseline LC window, based on the time lag between LC activity and pupil response (Yang et al., 2021).

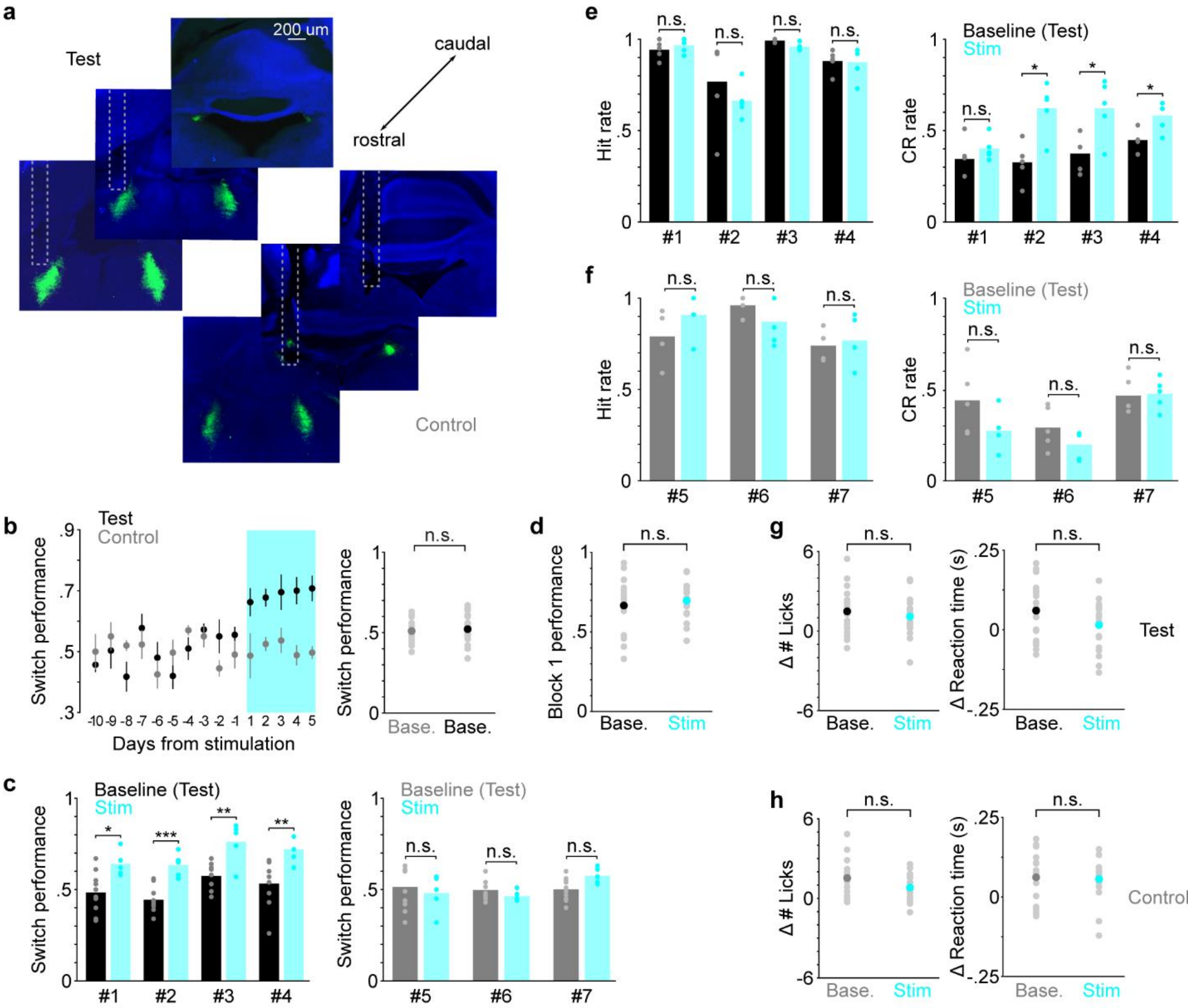


**Figure S3.**

**(a)** Three histological sections showing the LC (green) and the optical fiber tract from a test (Top, LC stimulation) and a control (Bottom, off-LC stimulation) mouse, respectively.

**(b)** Left: Group average switch performance for the test and control groups during an extended baseline period (10 consecutive days prior to stimulation) and optical stimulation period (5 consecutive days, cyan). Day -1 represents the last day without stimulation. Day 1 represents the first day with stimulation. Right: Switch performance during the 10-day baseline period for the test (n = 40 blocks) and control (n = 30 blocks) groups (Test vs. Control: 0.52 ± 0.02 vs. 0.51 ± 0.02, p = 0.62, rank sum = 255).

**(c)** Switch performance for individual mice (as in Fig. 3d, e), but compared between a 10-day baseline period and 5-day stimulation period (cyan) for the test (Left, n = 4; Baseline vs. Stimulation, Mouse #1: 0.48 vs. 0.64, p = 0.013; Mouse #2: 0.44 vs. 0.63, p = 0.001; Mouse #3: 0.58 vs. 0.76, p = 0.008; Mouse #4: 0.53 vs. 0.72, p = 0.0060, Permutation test) and control (Right, n = 3; Mouse #5: 0.52 vs. 0.48, p = 0.57; Mouse #6: 0.50 vs. 0.46, p = 0.31; Mouse #7: 0.50 vs. 0.58, p = 0.060, Permutation test) groups.

**(d)** Comparison of task performance in block 1 between Baseline (n = 20) and Stimulation (n = 20) sessions for the test group (Baseline vs. Stimulation: 0.67 ± 0.03 vs. 0.70 ± 0.02, p = 0.54, rank sum = 387).

**(e)** Hit rate (Left) and correct rejection rate (Right) for individual mice in the test group (n = 4). Hit rate, Baseline vs. Stimulation, Mouse #1: 0.94 vs. 0.97, p = 0.45; Mouse #2: 0.77 vs. 0.66, p = 0.44; Mouse #3: 0.99 vs. 0.96, p = 0.49; Mouse #4: 0.88 vs. 0.87, p = 0.90, Permutation test; Correct rejection rate, Baseline vs. Stimulation, Mouse #1: 0.34 vs. 0.40, p = 0.32; Mouse #2: 0.32 vs. 0.62, p = 0.023; Mouse #3: 0.37 vs. 0.62, p = 0.036; Mouse #4: 0.45 vs. 0.58, p = 0.024, Permutation test.
**(f)** Same as in (e) but for the control group (n = 3). Hit rate, Baseline vs. Stimulation, Mouse #5: 0.79 vs. 0.81, p = 0.21; Mouse #6: 0.96 vs. 0.87, p = 0.18; Mouse #7: 0.74 vs. 0.77, p = 0.71, Permutation test; Correct rejection rate, Baseline vs. Stimulation, Mouse #5: 0.44 vs. 0.27, p = 0.10; Mouse #6: 0.29 vs. 0.20, p = 0.13; Mouse #7: 0.47 vs. 0.48, p = 0.88, Permutation test.
**(g)** Comparison of changes in number of licks (Left, Δ# Licks, Before - After) and reaction time (Right, ΔRT) between baseline (n = 20) and stimulation (n = 20) sessions for the test group. Δ# Licks, Baseline vs. Stimulation: 1.48 ± 0.37 vs. 1.10 ± 0.33, p = 0.49, rank sum = 425; ΔRT, Baseline vs. Stimulation: 0.06 ± 0.02 vs. 0.01 ± 0.02 s, p = 0.13, rank sum = 455. For stimulation sessions, the change (Δ) was calculated by subtracting the variable (# licks or RT) quantified in hit trials during optical stimulation trials from the variable quantified in hit trials during the last 50 trials of the previous block (i.e., no optical stimulation). For baseline sessions, no optical stimulation was delivered, and the change (Δ) was calculated by subtracting the variable quantified in hit trials during the first 50 trials following the rule change from the variable quantified in hit trials during the last 50 trials of the previous block.
**(h)** Same as in (g) but for the control group. Left: Changes in number of licks, Baseline vs. Stimulation: 1.53 ± 0.37 vs. 0.81 ± 0.28, p = 0.17, rank sum = 266. Right: Changes in reaction time, Baseline vs. Stimulation: 0.06 ± 0.02 vs. 0.06 ± 0.02 s, p = 0.80, rank sum = 239.

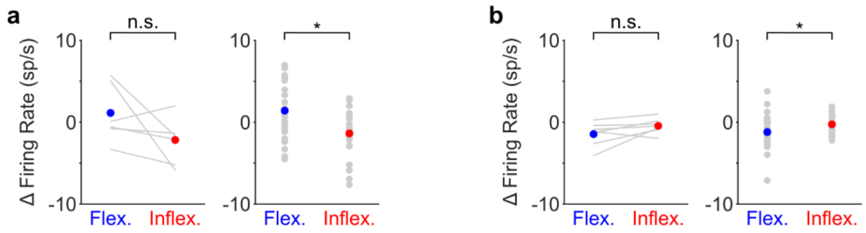


**Figure S4.**
**(a)** Changes in LC responses to whisker stimulation (Before - After) in hit trials during flexible and inflexible blocks (Left: paired Flexible vs. Inflexible: 1.14 ± 1.24 vs. -2.16 ± 1.01 spikes/s, p = 0.14, t-stat = 1.75, n = 6, two-tailed t-test; Right: unpaired Flexible vs. Inflexible: 1.41 ± 0.63 vs. -1.37 ± 0.66 spikes/s, p = 0.012, rank sum = 783, n = 47). LC responses to whisker stimulation were quantified in a 100-ms window beginning at stimulus onset, subtracted from LC activity quantified in a 0.5-s baseline window ending at stimulus onset. There were no hit trials during the Before period for 2 blocks from the original full dataset (n = 49), and they were removed from this analysis. 1 block was included in the original paired analysis (n = 7), so the associated flexible-inflexible pair was removed from this analysis.
**(b)** The same as in (a) but for LC responses to the auditory tone (Left: paired Flexible vs. Inflexible: -1.45 ± 0.55 vs. -0.44 ± 0.34, p = 0.11 spikes/s, signed rank = 4, n = 7; Right: unpaired Flexible (n = 28) vs. Inflexible (n = 21): -1.21 ± 0.38 vs. -0.26 ± 0.27 spikes/s, p = 0.042, rank sum = 599, n = 49). Tone responses were quantified in a 300-ms window beginning at tone onset, subtracted from LC activity quantified in a 0.5-s baseline window ending at tone onset.
